# Predicting bacterial-mediated entomopathogenicity through comparative genomics and statistical modeling

**DOI:** 10.1101/2025.03.24.645118

**Authors:** Daniela Yanez Ortuno, Melissa Y. Chen, Keegan McDonald, Allison Gacad, Juli Carrillo, Cara H. Haney

## Abstract

Bacterial genomes encode for vast functional diversity and have both beneficial and detrimental effects on insect hosts. However, connecting bacterial genotype to a host-associated phenotype can be experimentally time-consuming, particularly if insecticidal mechanisms are highly host-specific. To streamline the identification of virulence genes on a new host, we tested if merging existing mechanistic knowledge with *in vivo* tests on a small number of bacterial isolates could predict bacterial genes associated with entomopathogenesis. We used a model consisting of *Drosophila melanogaster* interactions with pathogenic and commensal genome-sequenced strains of *Pseudomonas* bacteria. We compiled a database of previously described insecticidal and biocontrol genes within the *Pseudomonas* genus and used comparative genomics to probe the distribution of these genes across *Pseudomonas* strains. We found natural variation in the presence of known insecticidal genes across the genus. We tested the insect-killing capacity of 13 *Pseudomonas* spp. strains against *D. melanogaster* and found natural variation in insecticidal activity. To identify bacterial genes associated with fly mortality, we employed two statistical models to correlate bacterial virulence with the presence of previously described insecticidal activity. To validate our predictions, we used a *P. aeruginosa* PAO1 transposon mutant library and identified 8 operons that are necessary for killing *D. melanogaster*. We show that by combining existing literature with phenotyping a small number of strains, we can identify genes necessary for insecticidal activity in a previously untested insect model. More broadly these findings illustrate a discovery pipeline for bacterial virulence mechanisms, accelerating the discovery of insect pest biocontrol mechanisms.

## Introduction

Insect pests significantly threaten global crop yields and food security. Pests account for 20–40% of global crop losses annually, a challenge further intensified by climate change and growing pesticide resistance [1]. For instance, many insect pests have developed insecticide resistance, sometimes to multiple insecticides with distinct modes of action, resulting in few existing treatments [2, 3]. Furthermore, climate change has resulted in insect range expansion and increased crop stress resulting in the need to rapidly develop new management tools [4–6]. These examples highlight the pressing need for new and rapid pest management strategies to target existing and emergent pests.

Bacteria represent a promising source of insecticidal compounds, offering a sustainable alternative to conventional pesticides [7–9]. Many bacterial species have evolved potent virulence mechanisms that enable them to kill insect hosts through diverse strategies, including toxin production, gut colonization, and immune suppression [10–12]. However, the efficacy of bacterial insecticides is influenced by whether they function as generalists, capable of infecting multiple insect species, or specialists, which exhibit narrow host specificity. Generalists, such as *Photorhabdus* Tc toxins, exert broad insecticidal effects through systemic toxicity mechanisms, including pore formation and immune modulation [10–14]. In contrast, specialists, such as *Bacillus thuringiensis* (Bt) Cry toxins, rely on highly specific receptor interactions in the insect midgut, making them effective but also susceptible to resistance via single-gene mutations [15, 16].

While many insecticidal mechanisms have been extensively studied, their predictability across different bacterial strains and insect hosts remains poorly understood. For instance, *Bacillus sphaericus* effectively targets *Culex* and *Anopheles* mosquitoes but is ineffective against *Aedes aegypti* due to receptor-based resistance mechanisms [16]. Similarly, bacterial colonization strategies, such as the Type VI Secretion System (T6SS), can play a crucial role in host interactions by modulating gut microbiota and delivering toxic effectors [18]. These observations highlight the need for systematic and scalable approaches to predict bacterial insecticidal activity across insect hosts.

In this study, we focused on *Pseudomonas* as a model bacterial system due to its genetic diversity, well-characterized virulence factors, and ability to colonize a wide range of environments and hosts[19–21]. *Pseudomonas* strains exhibit insecticidal properties through diverse virulence factors such as pore-forming toxins, proteases, and secondary metabolites [19–24]. *Drosophila melanogaster* serves as a valuable model for studying these interactions [25]. Key virulence determinants in *Pseudomonas* have been identified in *D. melanogaster* including Monalysin, a β-pore-forming toxin that damages the intestinal epithelium, and Fit toxins and cyclic lipopeptides like entolysin, which enhance pathogenicity under the control of the GacS/GacA regulatory system [26,27]. Despite this understanding of *Pseudomonas* virulence mechanisms, current methods for identifying bacterial insecticidal traits often rely on labour-intensive screening and mutant characterization, limiting the speed at which new candidates can be discovered.

To overcome these limitations, we developed a high-throughput, predictive framework that integrates existing literature with targeted experimental validation to systematically identify bacterial genes associated with insect killing. We constructed a database of insecticidal genes derived from literature on different bacteria and hosts and then we tested multiple *Pseudomonas* strains for their ability to kill *Drosophila melanogaster* and *Drosophila suzukii* via oral infection. Using two complementary statistical models, Cox Proportional Hazard and Random Forest Analysis, we correlated bacterial genetic profiles with insecticidal activity to identify genes predictive of virulence. This approach identified previously characterized genes known to be involved in *D. melanogaster* killing, as well as newly implicated genes from within our database. Collectively, this work establishes a method for the rapid identification of bacterial biocontrol agents and provides a framework for predicting bacterial insecticidal efficacy across diverse insect hosts, ultimately advancing pest management strategies.

## Material and Methods

### Bacterial strains and growth conditions

All the *Pseudomonas* strains and mutants used in this study are summarized in Table 1 and Table 2. Strains were streaked on LB agar and incubated at 28°C or 37° (*P. aeruginosa* only) for 24 hrs. Overnight cultures were made from single colonies, transferred to 100 mL of 3% NaCl LB, grown at 28 °C or 37 °C (*P. aeruginosa* only) and shaken at 180 rpm. PAO1 transposon insertion mutants were obtained from the PAO1 transposon insertion library [36]. Wildtype PAO1 and the transposon insertion mutants used in this study were cultured in LB. The initial screen (Figure 1B) was conducted using *Pseudomonas aeruginosa* PAO1 (H103), and while the parental strain of the transposon mutant library, *Pseudomonas aeruginosa* PAO1 (MPAO1), displayed slightly reduced killing activity compared to H103, as an additional control we chose to test neutral mutants from the library for comparison. Our analysis showed no significant difference in killing activity between the neutral mutants and the H103 strain (Figure S5).

**Figure 1.**
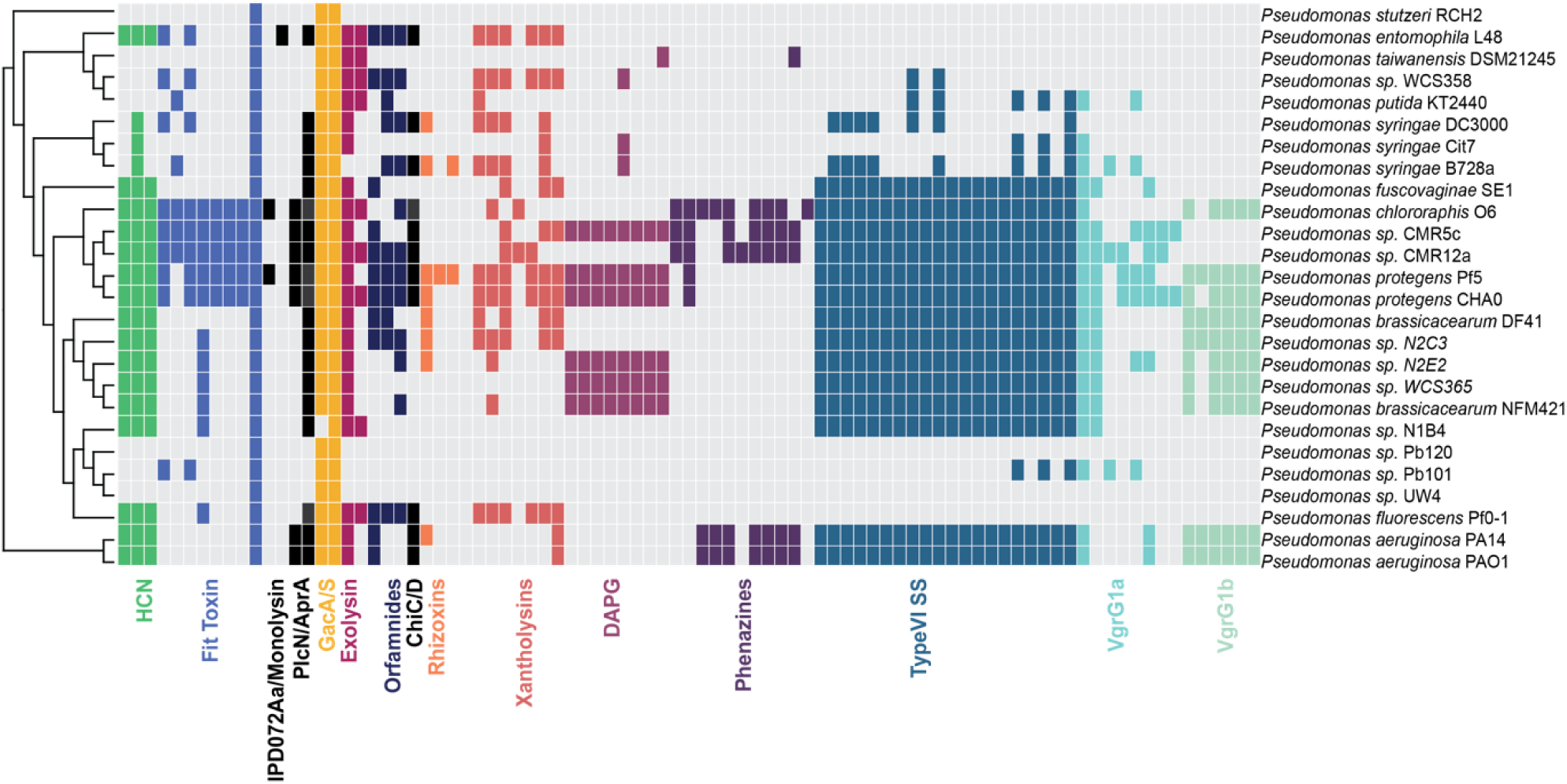
Distribution of insecticidal, biocontrol and antifungal genes within the *Pseudomonas* genus. Coloured squares represent the presence of homologous genes, while absence is represented by white. The genus *Pseudomonas* is divided into 5 phylogenetic groups, *Pseudomonas aeruginosa, P. fluorescens*, *Pseudomonas putida and Pseudomonas syringae.* Within the *P. fluorescens* group, we have other subgroups such as *P. chlororaphis* and *P. protegens.* The species tree was constructed using the PyParanoid comparative genomics tool [28]. Squares represent the presence and absence of individual genes associated with each locus based on PyParanoid presence-absence data.

**Table 1:** *Pseudomonas* strains used for the experiments in this study.

| Strain | Group | Subgroup | Number of genes | Oral Insecticidal Activity | Pathogenic to Plants |
| --- | --- | --- | --- | --- | --- |
| DSM21245 | *P. putida* | *P. putida* | 75 | This study | ---- |
| L48 | *P. putida* | *P. putida* | 120 | This study | No |
| cit7 | *P. syringae* | *P. syringae* | 75 | This study | No |
| B728a | *P. syringae* | *P. syringae* | 103 | This study | Yes |
| O6 | *P. fluorescens* | *P. chlororaphis* | 163 | This study | No |
| CMR5c | *P. fluorescens* | *P. protegens* | 213 | This study | No |
| CMR12a | *P. fluorescens* | *P. protegens* | 147 | This study | No |
| Pf5 | *P. fluorescens* | *P. protegens* | 240 | This study | No |
| CHA0 | *P. fluorescens* | *P. protegens* | 229 | This study | No |
| WCS365 | *P. fluorescens* | *P. fluorescens* | 143 | This study | No |
| N2E2 | *P. fluorescens* | *P. fluorescens* | 155 | This study | No |
| PAO1 | *P. aeruginosa* | *P. aeruginosa* | 137 | This study | Yes |
| PA14 | *P. aeruginosa* | *P. aeruginosa* | 139 | This study | Yes |

**Table 2:**
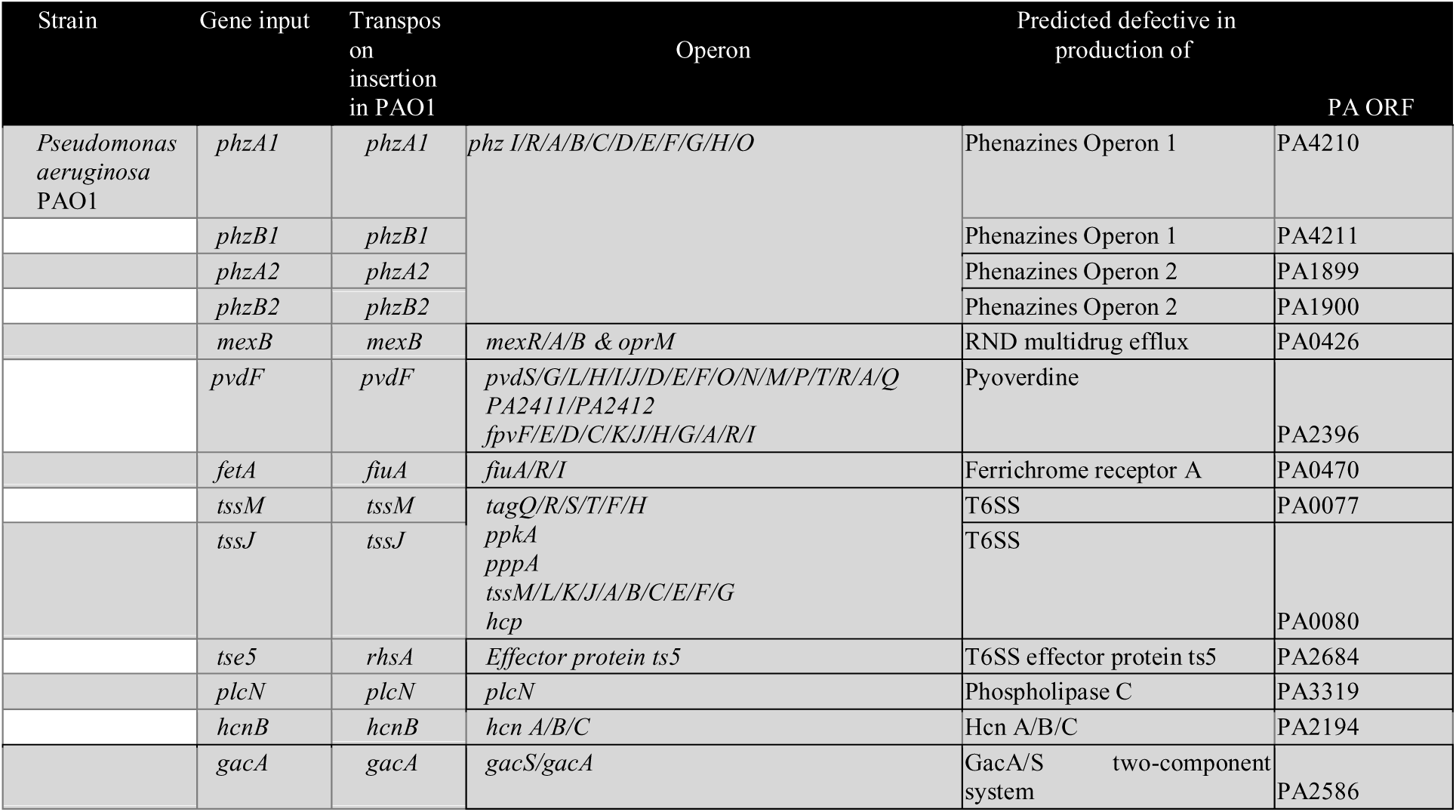
Mutant genotypes in *P. aeruginosa* PAO1 used in this study.

### Bacteria preparation for *D. melanogaster* and *D. suzukii* oral infections

Bacteria for infecting flies were prepared as described [25]. Freezer stocks were streaked onto LB-milk plates and incubated the plate at 28°C for 2 days. Colonies were selected and inoculated into 100 mL cultures in 3% NaCl LB broth and shaking at 180rpm for at least 20 hours at 28°C. Subsequently, cells were pelleted by centrifugation at 2,500 x g for 30 min at 4°C. After pelleting, the supernatant was centrifuged again to confirm the removal of most bacteria. The culture was then concentrated by resuspending the pellet in 1mL of supernatant. The OD_600_ was measured against a 3% NaCl LB broth blank using 1mL of liquid per cuvette, using 1 in 100 dilutions of the bacterial concentrates.

### D. melanogaster and D. suzukii growth conditions

The *D. melanogaster* Oregon R colony was grown for 16 hours of light and 8 hours of dark at 24.5 °C and 22.5°C respectively, with 50% humidity. Flies for the initial screen Figure 2A, were reared in freshly made Lewis medium. Flies for validation experiments (Figure 5) were reared in Bloomington media. *D. suzukii* was reared in milk bottles containing a standard potato flake diet (Ward’s® Instant Drosophila Medium) [25]. These bottles were incubated under a 16 h light:8 h dark at 24.5 °C and 22.5°C.

**Figure 2.**
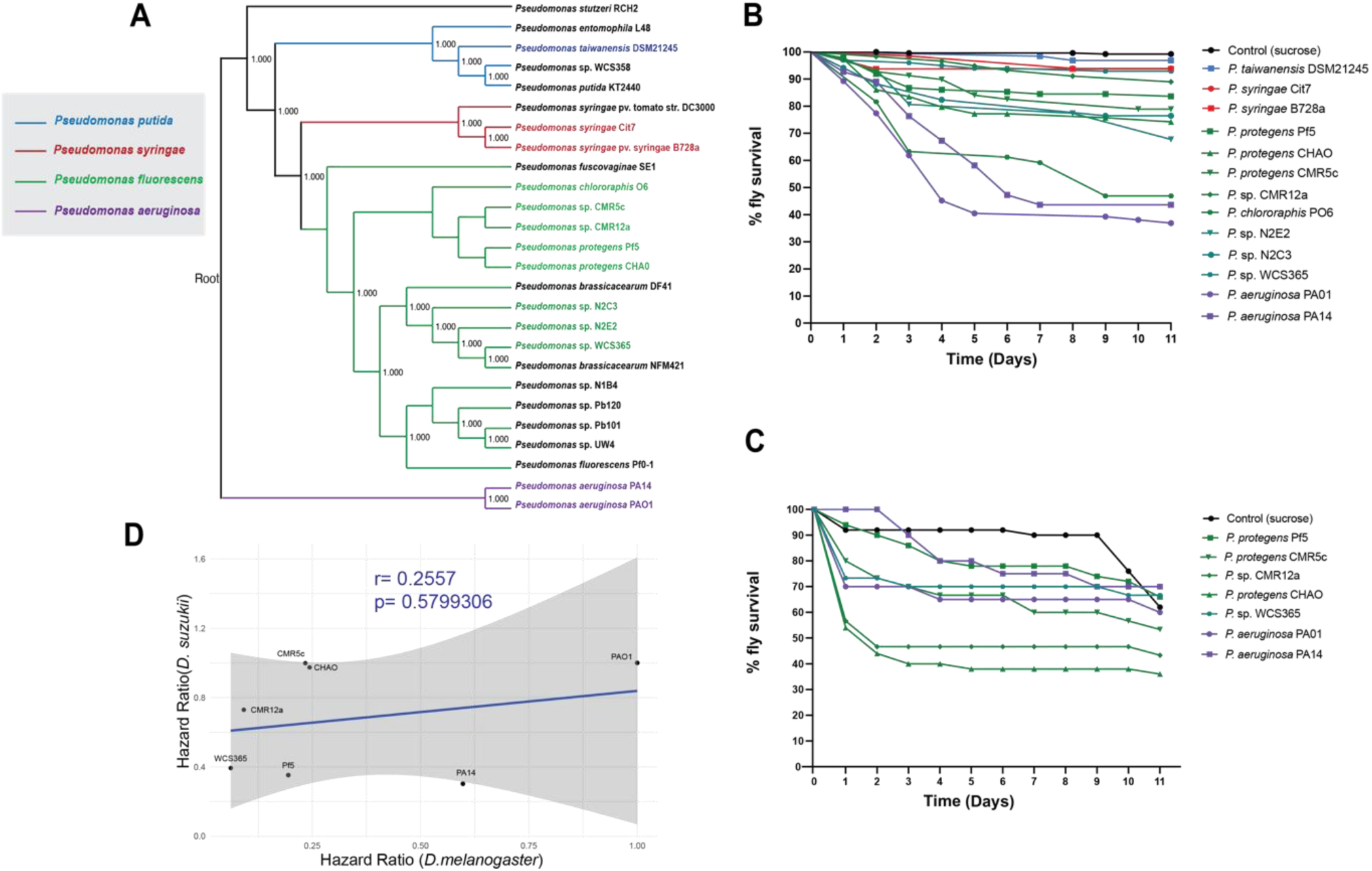
*Drosophila melanogaster* survival after oral infection with diverse *Pseudomonas* strains. A) Phylogenetic tree of *Pseudomonas* strains tested in the fly survival assay. Colours indicate different *Pseudomonas* species: *P. putida* (blue), *P. syringae* (red), *P. fluorescens* (green), and *P. aeruginosa* (purple). B) Kaplan-Meier (KM) survival curves of *D. melanogaster* Oregon-R flies following oral infection with *Pseudomonas* spp. strains Pf-5, CHA0, CMR5c, CMR12a, L48, cit7, WCS365, DSM21245, B728a, PO6, PA14, PA01 (OD600 = 100) or control 5% sucrose. The KM survival curve was calculated from 3 vials of 10 flies per treatment group. C) Kaplan-Meier (KM) survival curves of *D. melanogaster* Oregon-R flies following oral infection with *Pseudomonas* strain Pf5, CHA0, CMR5c, CMR12a, WCS365, PA14 and PA01 (OD600 = 100) or control 5% sucrose solution D) Linear regression on *D. melanogaster* and *D. suzukii* hazard ratios per strain. The shaded region represents the 95% confidence interval for the linear regression (blue line) with Pearson coefficient (r) and p-value.

**Figure 3.**
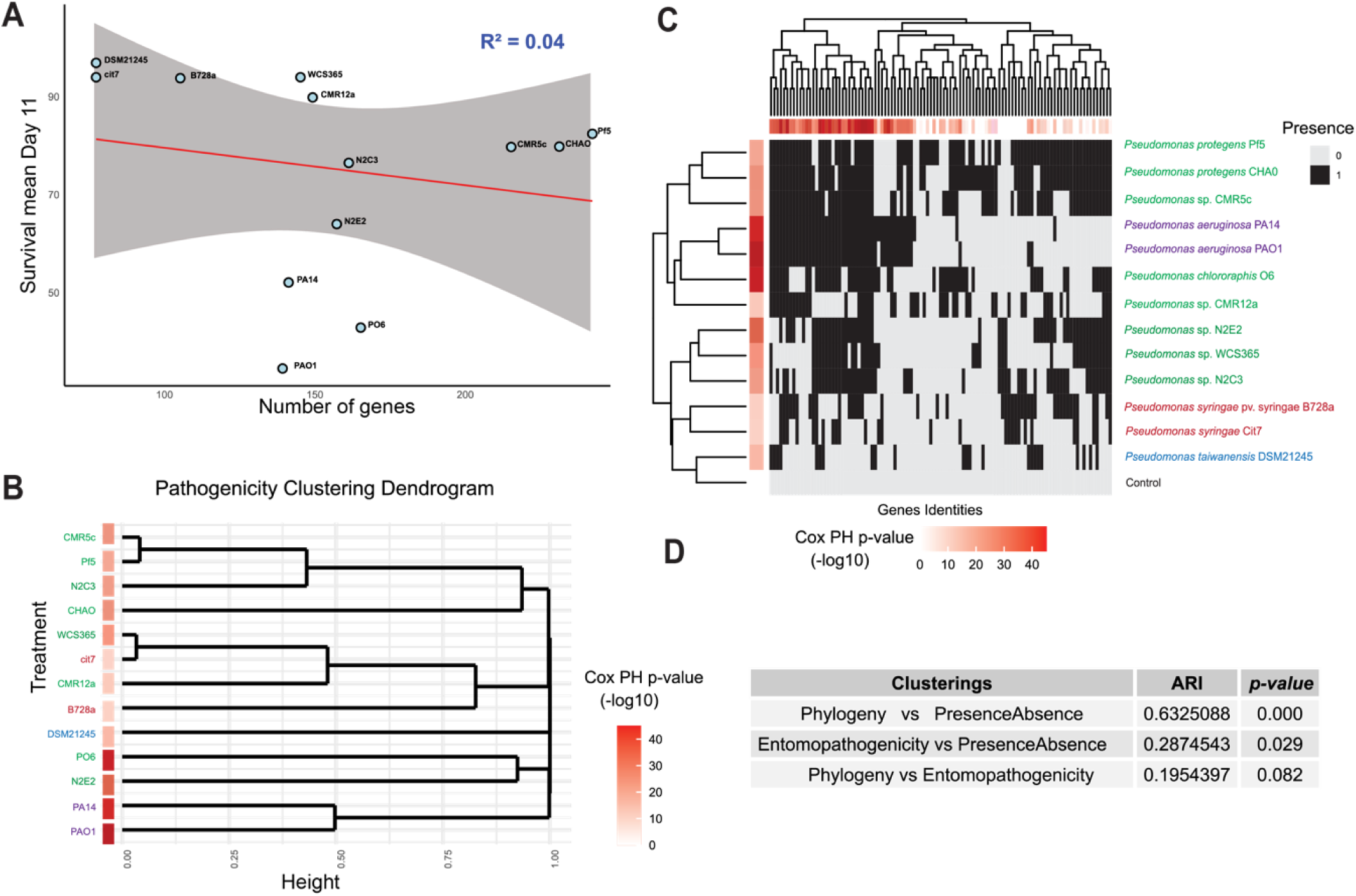
Presence of insecticidal and biocontrol genes, but not phylogeny, predicts insecticidal activity within the genus *Pseudomonas*. A) Linear regression on *D. melanogaster* survival on day 11 and the number of predicted insecticidal genes in the given strain. The shaded region represents the 95% confidence interval for the linear regression (red line). B) Clustering of bacterial strains based on *Drosophila* killing activity. C) Clustering between genes and strains based on presence/absence data. The top dendrogram shows genes clustered by their similarity across strains, while the side dendrogram shows strains clustered based on the similarity of their gene profiles. The horizontal red bars above the heatmap represent the p-values for how significantly each gene correlates with killing ability. The vertical red bars on the left side of the heatmap represent each strain’s proportional hazard (i.e., killing ability). D) Adjusted Rand Index (ARI) and p-values for the clustering analyses, comparing phylogeny, gene presence/absence, and entomopathogenicity.

### D. melanogaster and D. suzukii oral infection assay

Inoculum preparation, bacteria were concentrated to an OD600 of 100 in 5% sterile sucrose. To prepare flies, 2-5 days-old adult flies were collected and starved for at least 2 hours before the infection. Using forceps, filter paper discs were placed on top of the fly food, and 220 µL of inoculum was pipetted directly onto the filter paper. Once all vials were inoculated, we waited for 10 minutes for the inoculum to absorb into the filter paper before transferring flies into the infection tubes. Tubes were then incubated at 25°C and fly deaths within 2 hours were recorded, which were considered infection-independent deaths. The oral infection process was allowed to proceed for 24 hours, during which fly deaths were recorded. Following this initial 24-hour period, the flies were transferred to a fresh food source, and subsequent fly deaths were recorded daily over 10 days. Fly oral infection was repeated 3 times for each bacterial strain with 20 flies per replicate.

### Clustering analysis

Clustering similarity between phylogeny, gene presence/absence, and pathogenicity was evaluated using the Adjusted Rand Index (ARI), which measures agreement between clustering solutions while adjusting for chance. Observed ARIs were calculated for each pair of cluster labels, and their significance was assessed using 1,000 random permutations to generate null distributions. P-values were determined as the proportion of random ARIs greater than or equal to the observed values.

### Statistical curve analysis

Survival data in the *D. melanogaster* oral infection survival assays were analyzed using Prism 9 software using the Log-Rank test of the Survival package and the one-way ANOVA test. The hazard ratio for each of the strains compared to the control group was calculated using the Cox proportional hazard model in R (2022.2.0.443).

### Identification of insecticidal and biocontrol genes from literature

A list of bacterial genes reported to be involved in insecticidal activity towards different insects and the biocontrol of microbial pests was compiled. For each of the genes, amino acid sequences were collected from the NCBI gene bank and the *Pseudomonas* Genome Database. A database of 302 genes ranging from single genes to complete operons that were included in the analysis (Table S1).

### Bioinformatic analysis

#### Phylogenetic Tree Construction

A phylogenetic tree of bacterial strains was constructed using protein sequences extracted from each genome. The standalone OMA (Orthologous Matrix) algorithm [40], was employed with recommended parameter settings for the analysis. OMA groups were filtered using the filter_groups.py script [41], applying a minimum species coverage threshold of 10 out of 27 genomes. Orthologous groups were aligned with MAFFT (v7.526), allowing a maximum of 1000 iterative refinements. The resulting alignments were concatenated using the concat_alignments.py script [41]. Finally, a phylogenetic tree was generated using FastTree (v2.1.11).

#### Plotting presence and absence

Using PyParanoid, a pipeline for the rapid identification of orthologous genes in a set of genomes [28], and FastTree 2 a tool for inferring approximate-maximum-likelihood phylogenetic trees, a *Pseudomonas* species tree including representative strains from different clades was constructed (Figure 1). Using Pyparanoid, we identified orthologous groups in the *Pseudomonas* strains and plotted the presence and absence of these insecticidal-associated genes against the species tree (Supplementary Figure 1).

### Statistical models

#### Gene Annotation Dataset Preparation

The dataset consisted of bacterial gene metadata, including Gene ID, Gene Annotation, and Function. Gene ID represents the gene’s identity (e.g., *Fittoxin*), Gene Annotation describes genetic elements within the genome (e.g., *FitA, FitB*), and Function categorizes the gene based on literature into “Insecticidal,” “Biocontrol,” “Antimicrobial,” or various “Virulence factor” subtypes. The dataset was preprocessed to remove: 1) Gene IDs and annotations are present in all or no treatments. 2) Gene IDs associated with only one treatment, as their effects could not be distinguished from strain-level effects. 3) Gene IDs with identical occurrence patterns, which were collapsed into a single concatenated ID.

#### Cox Proportional Hazard Models (Model 1)

To evaluate fly mortality, Cox Proportional Hazard (Cox PH) models were applied. Fly mortality risk for each bacterial treatment was assessed using the coxph function in R’s *survival* package. Hazard ratio plots were created using the gg-forest function from the survminer R package[42]. Models included sex as a predictor, with females as the reference, and data were censored at 11 days post-exposure. Hazard ratios and BH-adjusted p-values for each gene were visualized in volcano plots (Figure S3) to identify genes significantly associated with fly mortality.

#### Random Forest Model (Model 2)

A random forest model was constructed using the *randomForestSRC* R package to evaluate the collective effects of gene IDs on fly mortality. This approach complemented the individual Cox PH models by accounting for interactions and co-occurrence patterns among genes, mitigating false positives.

#### Integrated Ranking Analysis

By combining Cox PH (p-value and coeff) and random forest (importance value) results into a ranking system (Figure 4D), genes were classified as insecticidal candidates by showing significance in both models.

**Figure 4.**
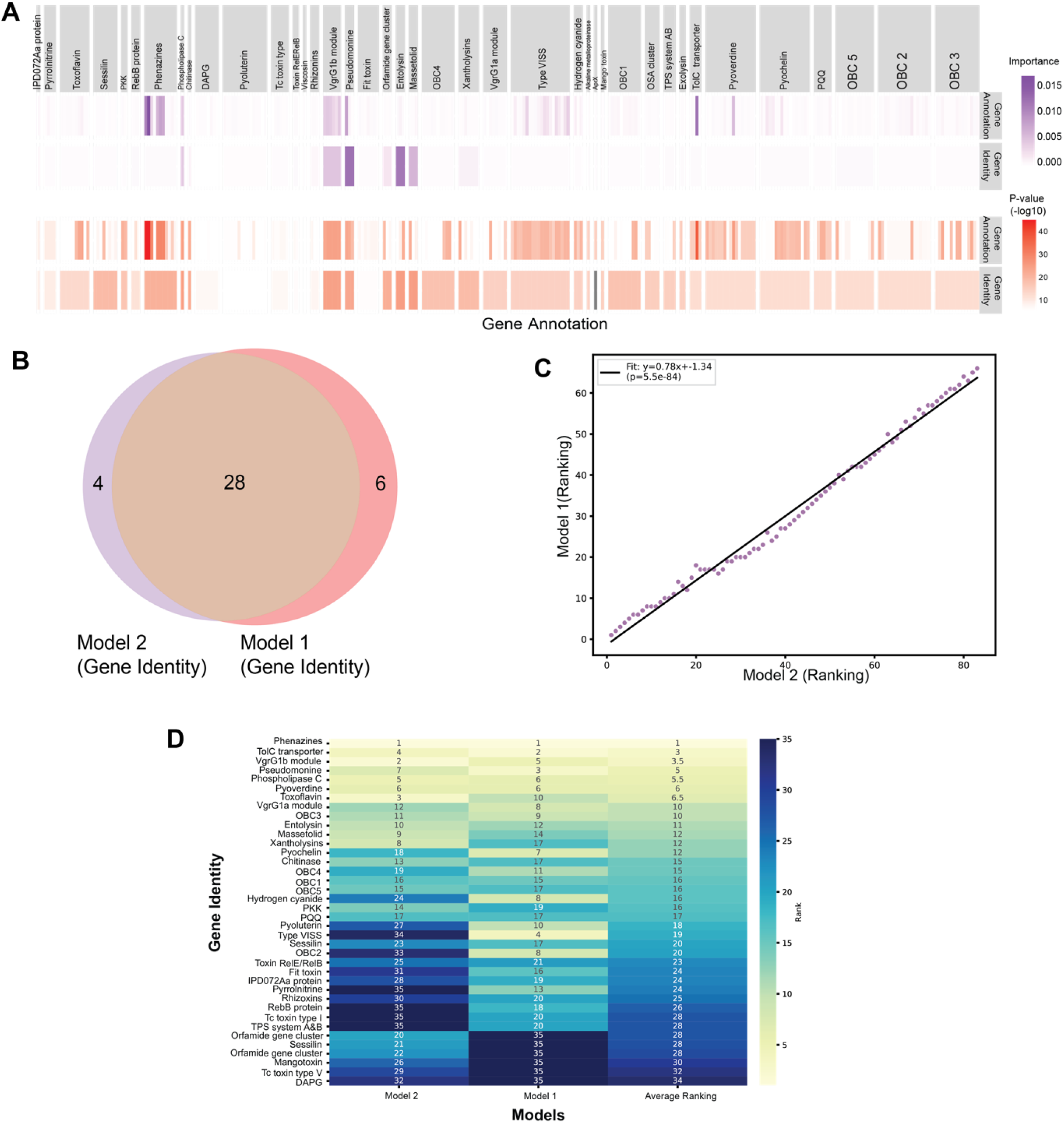
A subset of insecticidal and biocontrol genes predict virulence against *D. melanogaster*. A) Statistical models: Cox proportional hazard modeling and random forest analysis. The purple boxes on the top horizontal row show the individual gene identity risk *p*-value using the random forest analysis. The red boxes on the top horizontal row show the individual gene identity risk *p*-value using the Cox proportional hazard on individual genes. B) Venn diagram of gene identities for each model. Central overlap: shared genes. Unique segments: genes unique to each model (red segment for Model 1, purple segment for Model 2). C) Positive correlation between gene identity rankings in Model 1 (y-axis) and Model 2 (x-axis) (*p* < 0.05). C) Venn diagram of gene identities for each model. Central overlap: shared genes. Unique segments: genes unique to each model (red segment for Model 1, purple segment for Model 2). D) Heatmap of the ranking of gene identities for each model. The x-axis shows the gene identities and the y-axis the different rankings. Model 1 ranking is based on p-value and coefficient, Model 2 ranking is based on the importance value > 1 and average ranking is a combination of both rankings.

**Figure 5:**
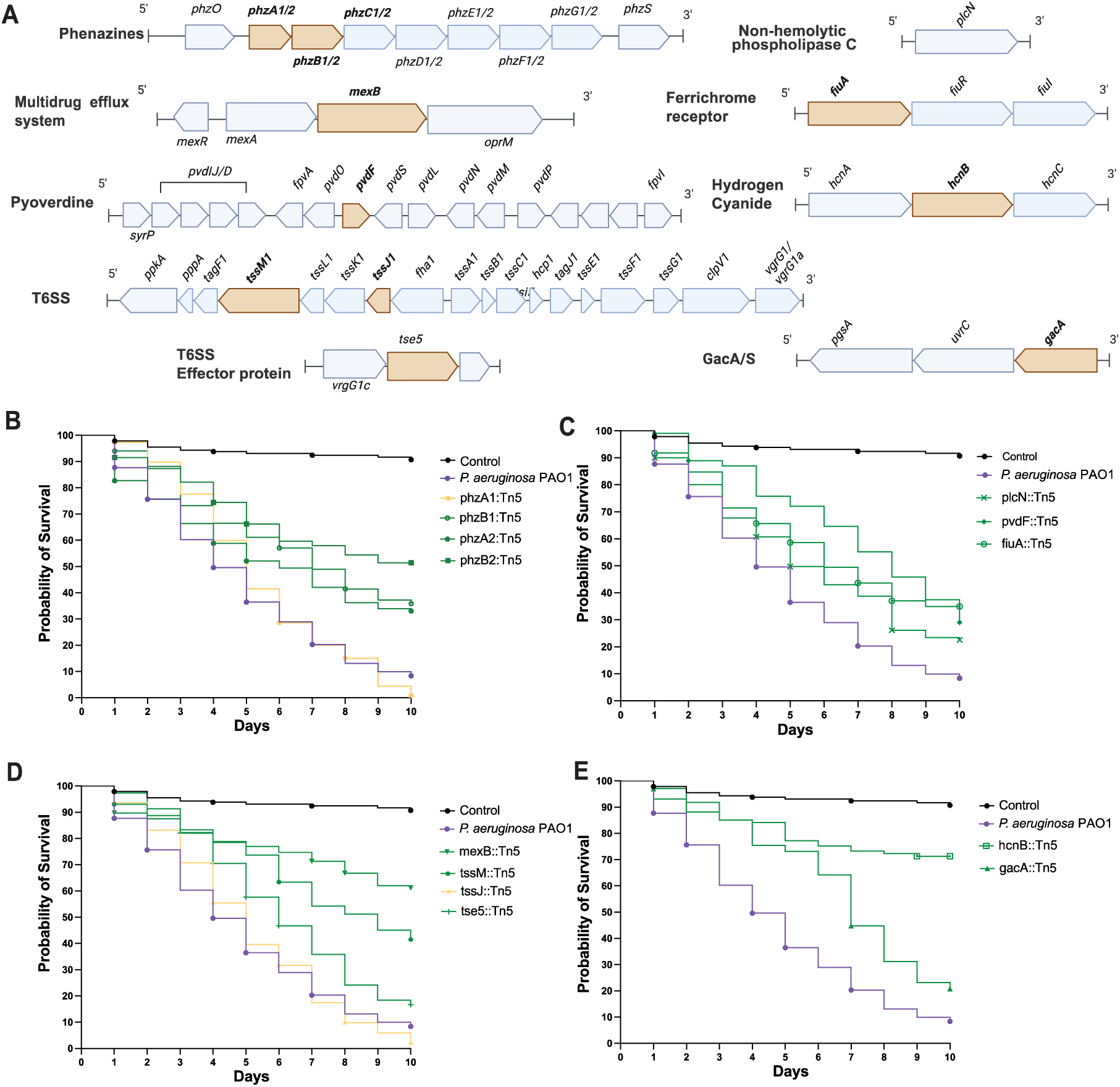
Modeling approaches identified *Pseudomonas* genes that are necessary for insecticidal activity in *Drosophila*. Genes predicted by both models (Figure 4) were validated using *P. aeruginosa* PA01 (H103) and PAO1 transposon mutants. A) Schematic representation of the seven distinct genes and operons selected from the models; the gene in tan indicates the location of the transposon insertion. B) Kaplan-Meier (KM) survival curves of *D. melanogaster* Oregon- R flies following oral infection with *Pseudomonas aeruginosa* PAO1 and transposon insertion mutants in genes encoding phenazine biosynthesis (*phz1 and phz2*); C) TolC transporter (*mexB*), and the Type VII secretion system (*tssM* and *tssJ*), including module VgrG1b (*tse5*); D) PlcN, ferrichrome receptor A (*fiuA*) and pyoverdine biosynthesis; E) GacA/S two-component system and hydrogen cyanide.

## Results

### Natural variation of insecticidal and biocontrol genes in *Pseudomonas*

To investigate the distribution of insect-killing activity of *Pseudomonas* towards *D. melanogaster,* we first identified previously described insecticidal and biocontrol genes from the literature and used comparative genomics to map their distribution across the genus *Pseudomonas.* Then, we tested the functional distribution of insecticidal activity by testing the killing activity of a subset of strains in *D. melanogaster* infection assays.

To assess the conservation of previously described insecticidal genes within the *Pseudomonas* genus, we conducted a comprehensive literature review and identified 302 genes previously implicated in insecticidal activity, virulence, or biocontrol against eukaryotic or prokaryotic organisms including insects, bacteria, and fungi. Using a comparative genomics database [28], we mapped the presence and absence of insecticidal and biocontrol genes onto a phylogenetic tree and visualized the distribution of these genes across different *Pseudomonas* groups (Figure 1). This analysis revealed variability in insecticidal and biocontrol gene presence between phylogenetic groups of *Pseudomonas*. Specifically, strains from the closely related subgroups *P. protegens* and *P. chlororaphis* have the highest number of insecticidal and biocontrol genes, with approximately 200 and 160 genes present, respectively from a total database of 302 query genes (Figure 1; Tables 1 and S1). As a result, we further investigated the distribution of insecticidal genes within the *P. protegens* and *P. chlororaphis* subgroups (Figure S2). Notably, the *fit* insect toxin gene cluster (*fitABCDEFGH*) and genes encoding the novel protein IPD072Aa were found to be uniquely associated with *P. protegens* and *P. chlororaphis*. IPD072Aa protects against Coleopteran pests when expressed in genetically modified plants [29] and was only found to be present in *P.* sp. WCS374, *P. protegens* Pf5, and *P. chlororaphis* O6 (Figure1 and Figure S2). These results suggest that some insecticidal mechanisms may be unique to subgroups of *Pseudomonas*.

We also observed *Pseudomonas* subgroup-specific distributions of additional previously described insecticidal and biocontrol genes. The phenazine cluster, responsible for the production of the antimicrobial compound Phenazine-1-carboxylic acid (PCA) was conserved in *P. aeruginosa* and *P. chlororaphis* but was absent in all other *Pseudomonas* groups including *P. protegens* (Figure 1). Conversely, genes encoding 2,4-Diacetylphloroglucinol (DAPG) biosynthetic enzymes, a known antifungal biocontrol molecule, were exclusive to *P. fluorescens* and *P. protegens* (Figure 1 and Figure S2). Interestingly, genes encoding the high-molecular-weight Tc toxin (Tcc) were distributed across the *P. fluorescens* group but absent in both *P. protegens* and *P. chlororaphis*. These findings suggest that insecticidal and biocontrol genes are differentially distributed across the *P. fluorescens* group, with most genes from Table 1 conserved in the *P. protegens* and *P. chlororaphis* subgroups. However, functional testing is required to elucidate the role of these gene distributions in insect-killing activity.

### Natural variation of *Pseudomonas* insecticidal activity in *Drosophila* spp

To determine which *Pseudomonas* strains, exhibit insecticidal activity against *D. melanogaster* (and the closely related agricultural pest *D. suzukii),* we orally infected flies with representative strains from the different *Pseudomonas* subgroups, and monitored their survival over 11 days (Figure 2). Kaplan-Meier survival analysis, including log-rank tests and pairwise comparisons, revealed a broad range of insecticidal activity among *Pseudomonas* strains. *P. putida* (DSM21256) and *P. syringae* (cit7 and B728a) exhibited minimal to no insecticidal activity, with fly survival rates of 95–100%. Similarly, *P. fluorescens* group strains (WCS365, N2E2, and N2C3) showed moderate pathogenicity, with survival rates ranging from 70% to 90% (Figure 2B). The most potent insecticidal strains were *P. aeruginosa* and *P. chlororaphis*, with significantly lower survival rates. Flies infected with *P. aeruginosa* PA14 and PAO1 had survival rates of 45% and 40%, respectively, while *P. chlororaphis* PO6 resulted in 50% survival (Figure 2B). These findings highlight significant natural variation in oral insecticidal activity within the *Pseudomonas* genus against *D. melanogaster*.

To quantitively differentiate the killing profiles of *Pseudomonas* strains against *D. melanogaster*, we applied a Cox proportional hazards model (Figure S3A) to characterize the strains based on their killing efficiency. The Cox model categorized *Pseudomonas* strains into distinct groups: high killing (<20 hazard ratio and *p* < 0.001), medium killing (10–20 hazard ratio and *p* < 0.001), and low killing (1–10 hazard ratio and *p* < 0.002) profiles. The low-killing group included strains DSM21245, Cit7, WCS365, CMR12a and B728a, the medium-killing group consisted of CHAO, Pf5, CMR5c, N2C3 and the high-killing strains were N2E2, PO6 and PA14, and PAO1. These results suggest that *Pseudomonas* strains exhibit discrete killing profiles against *D. melanogaster*, potentially governed by shared genetic and molecular mechanisms.

*D. suzukii*, or the spotted wing *Drosophila*, is a major agricultural pest capable of laying eggs in ripening fruits, causing significant economic losses [30,31]. Based on our observation that some *Pseudomonas* strains are effective against *D. melanogaster* we hypothesized that some strains may also exhibit activity against *D. suzukii*. To test this hypothesis, we analyzed the activity of selected strains of *Pseudomonas fluorescens* (CHA0, Pf5, CMR12a, CMR5c, and WCS365) and *Pseudomonas aeruginosa* (PA14 and PAO1) against adult *D. suzukii*. Our findings revealed that most strains effective against *D. melanogaster* also affected *D. suzukii* (Figure 2C). Flies in the control group (sucrose) exhibited the highest survival until day 10, with minimal mortality observed throughout the experiment. In contrast, flies treated with *P. protegens* CHA0, *P. sp.* CMR12a*, P. sp.* CMR5c and *P. aeruginosa* PA01 displayed rapid declines in survival within the first few days, indicating high virulence. Other strains, such as *P.* sp. WCS365*, P. protegens* Pf5 and *P. aeruginosa* PA14, exhibited intermediate effects, with survival falling between the control and highly virulent strains (Figure 2C). These results indicate that while there is overlap in the killing profiles against these closely related fly species, they have distinct susceptibilities to some bacterial strains, in particular *P. aeruginosa* species.

To robustly test whether insecticidal activity against *D. melanogaster* would be predictive of killing *D. suzukii,* we performed a linear regression to test the association between the survival of each species. We found a weak, and not statistically significant, positive linear relationship between the hazard ratios of the strains tested in both insect models, as indicated by a Pearson correlation coefficient of 0.2557 (p = 0.580) (Figure 2D). A Spearman correlation analysis yielded a coefficient of 0.3214 (p = 0.498). This indicates that insecticidal activity against one species may not be broadly predictive, against even members of the same insect genus. Based on these findings, we cannot currently extrapolate the data from *D. melanogaster* to *D. suzukii* further emphasizing the need for novel approaches to identify insecticidal genes and traits. Moving forward, we decided to explore exclusively insecticidal activity in *D. melanogaster*.

### Cox proportional hazard modelling and random forest analysis predict entomopathogenic genes

We conducted a correlation analysis to explore the relationship between the insect-killing activity of bacterial strains and their genetic composition. Specifically, we examined the correlation between the percentage survival of *D. melanogaster* by day 11 and the number of insecticidal and biocontrol genes present in the strains we inoculated (Tables 1 and S1). The analysis revealed a weak negative correlation, as shown in Figure 3A, indicating that strains with more predicted insecticidal genes were associated with lower fly survival rates, suggesting increased killing activity. The R-squared value of 0.04 indicates that the total number of these genes does not fully explain the observed insecticidal activity (Figure 3A). Consistent with host-specific virulence mechanisms, this suggests that not all previously described biocontrol and insecticidal genes contribute to the insecticidal activity of *Pseudomonas* strains against *D. melanogaster*.

*Pseudomonas* virulence genes can be horizontally transferred over short evolutionary distances [32, 33]. At the same time, other traits correlate with phylogeny [34, 35], we hypothesized that insecticidal activity may be influenced by both phylogeny and gene presence/absence patterns, but to different extents. From Figure 1B, we observed that *P. aeruginosa* and *P. protegens* strains exhibited higher insecticidal activity than other *Pseudomonas* groups. To explore whether phylogeny explains this pattern, we clustered strains based on their killing profile (Figure 3B) and compared them to phylogeny. To test whether insecticidal activity and the presence/absence of known insecticidal genes correlate with phylogeny, we constructed a dendrogram based on gene presence/absence (Figure 3C). We quantified these relationships using the Adjusted Rand Index (ARI), which measures clustering similarity while accounting for random agreement (-1 = no agreement, 1 = perfect agreement). Pathogenicity showed a weak correlation with phylogeny (ARI = 0.1954, Figure 3D), suggesting phylogeny alone does not explain killing profiles. However, gene presence/absence clustering showed a slightly better correlation with pathogenicity (ARI = 0.2624), indicating that certain genes better predict insecticidal activity. The strongest correlation was between gene presence/absence and phylogeny (ARI = 0.6325), suggesting that while phylogeny shapes the distribution of insecticidal genes, specific genes may play a key role in pathogenicity.

We asked whether certain genes are better predictors of the killing of *D. melanogaster*. To address this, we employed two complementary strategies. In Model 1, we used a Cox proportional hazards model to assess each genetic predictor individually, iterating through genes from our comprehensive database. This method fitted separate Cox models for each predictor against entomopathogenicity (Figure 4A, S3A). Genes strongly associated with increased mortality (high hazard ratios and low p-values) were visualized using a volcano plot (Figure S4B). Using this model, we identified 34 distinct genes significantly correlated with entomopathogenicity (Figure 4B). To address potential false positives that may arise due to the correlation of insecticidal activity and strain phylogeny, we employed a dual-method approach that combined the Cox PH model with a Random Forest model (Model 2). This method evaluated all genes collectively, considering co-occurrence patterns and interdependencies. We ranked gene identities based on *p*-value and coefficient (Model 1) and importance value (Model 2), finding a strong positive correlation between the rankings, indicating that both models prioritized gene identities in a similar order (Figure 4C). The genes identified through this model (Figure 4A, purple panel) were largely a subset of those identified by the Cox model (Model 1). Model 2 identified 32 significant gene identities, including 28 overlapping with Model 1 and 4 unique genes to Model 2 (Figure 4B). The high degree of overlap from both models, which use independent approaches, suggests the 28 genes identified by both models are strong candidates for genes important for insecticidal activity against *D. melanogaster*.

### Modelling predicts genes necessary for insecticidal activity in *Drosophila melanogaster*

To determine if testing a limited number of strains could predict genes important for killing a new host, we tested mutants in genes predicted by our modelling approach (Figure 4). To validate the model, we made use of a previously described *P. aeruginosa* PAO1 transposon mutant library [36]. We chose 7 of the 28 genes to test by 1) for Model 1 ranking based on the *p*-value and coefficient (hazard ratio), which quantify the statistical significance and strength of a gene’s impact on fly mortality, and 2) for Model 2 genes were ranking genes normalized importance values from the Random Forest algorithm, which quantifies the contribution of each single gene to the overall predictive performance. We integrated findings from both models to identify high-ranking genes (Figure 4D). These genes included those involved in the production of phenazines, cyclic lipopeptides, and type VI secretion systems. We selected 7 functionally distinct genes or operons or individual genes for testing, prioritizing those involved in different processes including genes previously reported insecticidal or genes for biocontrol but not insecticidal activity (Table 2). These genes include phenazines, MexAB-OprM efflux pump, phospholipase C, the Type VII secretion system, pyoverdine, ferrichrome receptor A (fetA-like protein), hydrogen cyanide and the GacA/S two-component system (Figure 5A).

We found that of the 13 mutants tested, 11 had significant loss in virulence. These include transposon insertions in *phzB1, phzA2, phzB2, mexB, pvdF, fiuA, tssM, tse5, plcN, hcnB* and *gacA* (Figure 5). Disruption of phenazine production (*phzA2, phzB2,* and *phzB1*) significantly reduced virulence, though complete survival restoration may be hindered by redundancy between the two phenazine operons [35] (Figure 5B). Mutants in *mexB* (MexAB-OprM Efflux Pump) and *plcN* phospholipase C, and *pvdF* and *fiuA* (siderophore biosynthesis and iron acquisition) also demonstrated markedly reduced virulence (Figure 5 C, D). Loss of *tssM* and *tse5* impaired T6SS function, with the *tssM::*Tn5 mutant showing insecticidal activity reduction (Figure 5D). The *hcnB* mutant showed the highest virulence attenuation, underscoring the role of hydrogen cyanide in infection (Figure 5E). Surprisingly, the *gacA* mutant exhibited only a slight virulence reduction, suggesting a subtler role for this regulator (Figure 5E). Growth assays confirmed comparable growth rates among mutants and wild type when growing in minimal media supplemented with fly extract, ruling out generalized growth defects as the cause of virulence loss (Figure S6). These findings show that our models successfully identified genetic pathways required for the *Pseudomonas* pathogenicity of flies, indicating that comparative genomics coupled with testing a small number of strains successfully identified virulence factors in a new model.

## Discussion

In this manuscript, we explored the possibility of combining existing literature with functional tests from a limited number of strains to identify genes necessary for bacterial virulence in a specific insect model. We evaluated the genetic basis of insecticidal activity in target *Pseudomonas* strains against *Drosophila melanogaster* and, using robust statistical models, predicted which genes were crucial for killing activity against *D. melanogaster*. By integrating Cox proportional hazards modelling with Random Forest analysis, we developed a system capable of distinguishing key virulence and entomopathogenicity genetic determinants based on killing profile. This approach validated previously identified virulence genes from other models and identified novel genetic contributors, showcasing the power of these models in advancing our understanding of microbial entomopathogenicity.

The Cox proportional hazards model identified a broad set of genes significantly associated with reduced survival in *Drosophila melanogaster*, while Random Forest analysis highlighted a subset of these genes under stricter criteria, thereby minimizing the likelihood of false positives. The intersection of these models revealed 28 high-confidence gene candidates, including those involved in phenazine production, efflux pumps, siderophores, T6SS, and hydrogen cyanide synthesis. These findings emphasize the importance of using diverse statistical methods to enhance predictive accuracy.

The genes identified by both models underscore the complexity of insecticidal activity, where multiple genetic pathways converge to mediate virulence. For example, phenazine operons, previously associated with antimicrobial activity [37–39], emerged as significant determinants of insect mortality, suggesting a dual role in bacterial fitness and pathogenicity. Similarly, the MexAB-OprM efflux pump and some siderophores were shown to be critical for killing efficiency, likely by helping bacteria evade the insect immune barriers. These results highlight how statistical modelling can uncover genes with roles in virulence that might be overlooked in traditional single-gene studies.

We found a weak correlation (R² = 0.04) between the total number of insecticidal genes identified through a literature search and observed virulence, consistent with the observation that not all insecticidal mechanisms will be effective on a new host. This suggests that insecticidal activity against one species may not be broadly predictive, even within members of the same insect genus. Instead, our models demonstrate that specific genes better predict insecticidal profiles. The models developed in this study have broad applications beyond the insecticidal activity of the genus *Pseudomonas*. By prioritizing genes based on their statistical and functional significance, our approach can be adapted to other microbial systems to identify key determinants of pathogenicity or biocontrol potential. These insights also inform the design of biocontrol agents, enabling the selection of strains enriched with high-impact genes while minimizing unintended effects on non-target organisms. For example, our results reveal that different *Pseudomonas* strains exhibit varying efficacy in killing two closely related *Drosophila* species, with some strains being more effective against *D. suzukii* than *D. melanogaster*. This underscores the potential of our study to identify genes in *Pseudomonas* strains that target particular insect pests.

By using a combined approach, this study provides information on which genes to prioritize when screening insecticidal strains from rhizosphere communities. Using the results from our statistical model, it is possible to analyze new soil or environmental bacterial isolates to identify genes associated with the desired insecticidal phenotype. While the identified genes provide a strong starting point for understanding insecticidal phenotypes, their effectiveness against entirely new insect targets remains uncertain. Instead, the strength of this approach lies in its ability to guide the discovery of novel insecticidal traits tailored to agricultural applications. Future studies could explore whether the statistical models and identified genes can help predict insecticidal activity in diverse ecological contexts. This would involve validating the approach in systems with varying host-pathogen interactions and adapting it for targeted screening efforts to meet specific agricultural needs.

In conclusion, this study demonstrates the use of statistical models in identifying key genetic drivers of insecticidal activity, offering a framework for understanding microbial pathogenicity and its applications in biocontrol.

## Acknowledgements

This work was supported by and NSERC Discovery Grant and Accelerator Award (no. NSERC-RGPIN-2021-03587 to C. H.H. and Organic Science Cluster 3 (funding to C.H.H. and J.C.), led by the Organic Federation of Canada in collaboration with the Organic Agriculture Centre of Canada at Dalhousie University, supported by Agriculture and Agri-Food Canada’s Canadian Agricultural Partnership - AgriScience Program. Funding for D.Y.O. was supported through a UBC 4Y Fellowship and an NSERC-CREATE program. M.Y.C salary support was provided through an NSERC Banting Postdoctoral Fellowship (202209BPF-489437-BNE-CAAA-91388)

