## Supplementary material for "Predicting bacterial-mediated entomopathogenicity through comparative genomics and statistical modeling": Figure S1

### Supplementary Information

Supplemental Tables and Figures

Table of insecticidal genes (references and insects)

**Table S1. List of predicted insecticidal gene functions.**

| Identity | Gene Annotation | Source | Function |
|---|---|---|---|
| Hydrogen cyanide | *hcn A/B/C* | [1] | Insecticidal |
| Fit toxin | *fitA/B/C/D/E/F/G/H* | [2] | Insecticidal |
| IPD072Aa protein | *IPD072Aa* | [3] | Insecticidal |
| ß-Pore-Forming Toxin | *monolysin* | [4] | Insecticidal |
| Phospholipase C | *plcN* | [5] | Insecticidal |
| Alkaline metalloproteinase | *aprA* | [6] | Insecticidal |
| Two-component system | *gacS/gacA* | [7] | Insecticidal |
| Two-partner secretion toxin (Exolysin) | *ExlB/A* | [8] | Insecticidal |
| Orfamide gene cluster | *OfaA/B/C* | [9] | Insecticidal |
| Chitinase | *ChiC/D* | [7] | Insecticidal |
| Cyclic Lipopeptides (Rhizoxins) | *RzxB/C/D/E/F/G* | [10] | Biocontrol |
| Cyclic Lipopeptides (Xantholysins) | *ClpA/B/C/D/E/F/G/H* | [11] | Antimicrobial |
| 2,4-diacetylphloroglucinol (DAPG) | *PhlA/B/C/D/E/F/G* | [7] | Antimicrobial |
| Phenazines | *phzI/R/A/B/C/D/E/F/G/H/O glu* | [12] | Biocontrol |
| TPS System A&B | *tpsB1/A1/B2/A2/B3/A3/B4/A4* | [13] | Insecticidal |
| RebB protein | *rebB_1/reb_2/PPRCHA0_0184* | [14] | Insecticidal |
| OSA cluster | *PPRCHA0_4348-4354* | [13] | Insecticidal |
| O-antigenic polysaccharides (OBC1/OBC2/OBC3/OBC4/OBC5) | *OBC1 (PFL_2023-2033)*<br>*OBC2 (PFL_3077-3094)*<br>*OBC3 (PPRCHA0_1948-1966)*<br>*OBC4 (PFL_5482-5496)*<br>*OBC5 (PFL_5091-5108)* | [13] | Insecticidal |

| | | | |
|---|---|---|---|
| Polyphosphate Kinase | *pap* | [13] | Insecticidal |
| Type VISS | *tagQ/R/S/T/F/H*<br>*ppkA*<br>*pppA*<br>*TssM/L/K/J/A/B/C/E/F/G*<br>*hcp* | [15] | Biocontrol |
| VgrG1a&1b module | *vgrG1a & PPRCHA0_6003-6009*<br>*PPRCHAO_3010-3015* | [15] | Biocontrol |
| TolC transporter | *mexR/A/B & oprM* | [16] | Virulence factor |
| Pyoluterin | *pltL/M/R/A/B/C/D/E/G/Z/I/J/K/N/O/P/* | [17] | Antimicrobial |
| Pyrrolnitrine | *prnA/B/C/D* | [18] | Antimicrobial |
| Massetolid | *massA/B//C* | [19] | Antimicrobial |
| Sessillin | *sesA/B/C/T/R/D/B/C* | [9] | Antimicrobial |
| Metallopeptidase AprA | *aprX* | [7] | Insecticidal |
| Viscosin | *viscB/C* | [20] | Antimicrobial |
| PKK | *pkk1, pkk2A/B/C* | [21] | Virulence factor |
| Tc toxin | *tcaA1, tcaB1, tcdC1, tccC2_2, tccC2, Pfl01_0947* | [22] | Insecticidal |
| Toxoflavin | *toxH/G/M/R/E/C/B/D/A/F* | [23] | Antimicrobial |
| Pseudomonine | *pmsE/C/A/B* | [24] | Virulence factor (siderophores) |
| Pyochelin | *pchA/B/C/K/F/E/I/H/D/R*<br>*fetA/B/C/D/E/F & PFL_3504* | [25] | Virulence factor (siderophores) |
| Pyoverdine | *pvdS/G/L/H/I/J/D/E/F/O/N/M/P/T/R/A/Q*<br>*PA2411/PA2412*<br>*fpvF/E/D/C/K/J/H/G/A/R/I* | [26] | Virulence factor (siderophores) |
| PQQ | *gcd & pqqH/I/J/K/M/E/D/C/B/A/F* | [27] | Antimicrobial |
| Toxin RelEB | *relE/B* | [28] | Virulence factor |
| Entolysin | *eltA/B/C* | [29] | Insecticidal |
| Mangotoxin | *mgoA* | [30] | Virulence factor (plants) |

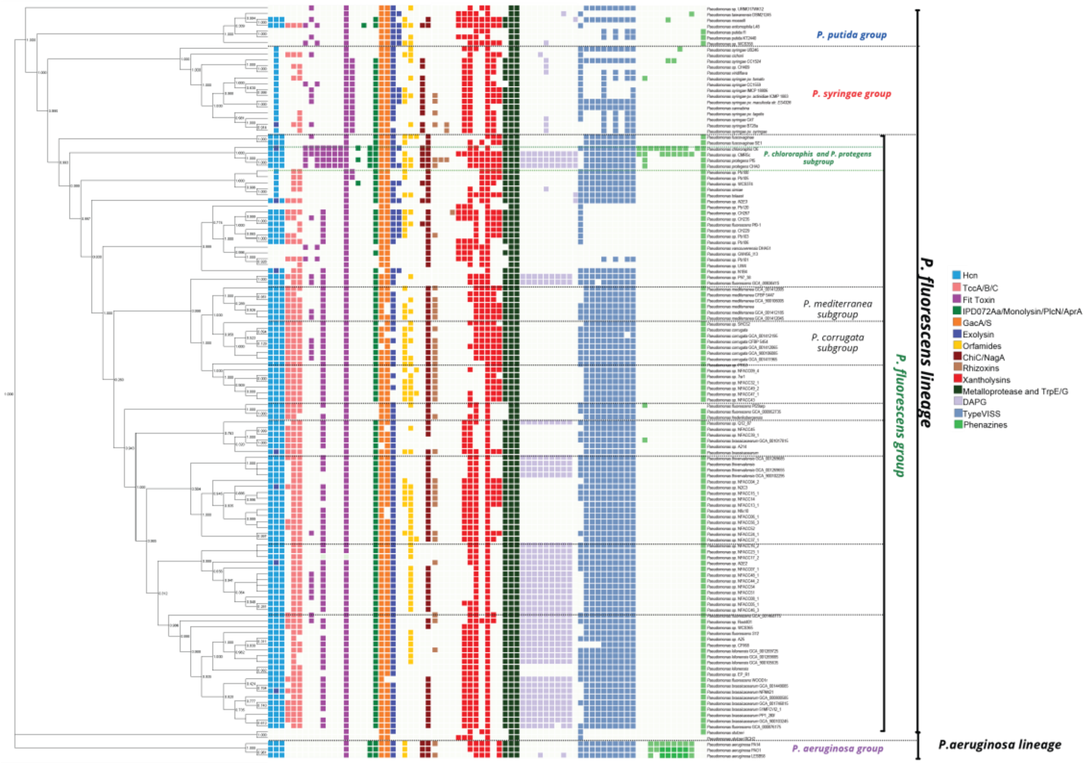


**Figure S1: Distribution of insecticidal, biocontrol and antifungal genes within the *Pseudomonas genus*.** The species tree was constructed using the PyParanoid comparative genomics tool. Squares represent the presence and absence of individual genes associated with each locus based on PyParanoid presence-absence data. Coloured squares represent the presence of a homologous gene, while absence is represented by white. The genus *Pseudomonas* is divided into 5 phylogenetic groups, *Pseudomonas aeruginosa*, *P. fluorescens*, *Pseudomonas putida and Pseudomonas syringae*. Within the *P. fluorescens* group, we have other subgroups such as *P. chlororaphis* and *P. protegens.*

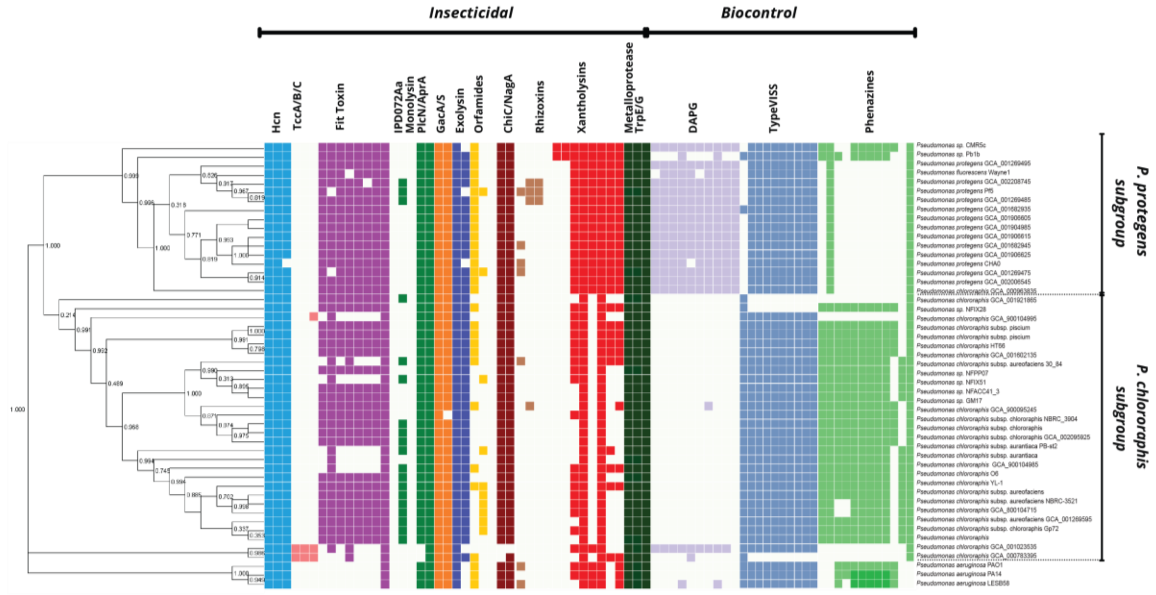


**Fig S2. Distribution of insecticidal, biocontrol and antifungal genes within the *Pseudomonas protegens* and *chlororaphis subgroup*.** The species tree was constructed using the PyParanoid comparative genomics tool. Squares represent the presence and absence of individual genes associated with each locus based on PyParanoid presence-absence data. Coloured squares represent the presence of a homologous gene, while absence is represented by white. The tree exclusively shows the *P. fluorescens* subgroups *P. protegens* and *P. chlororaphis.*

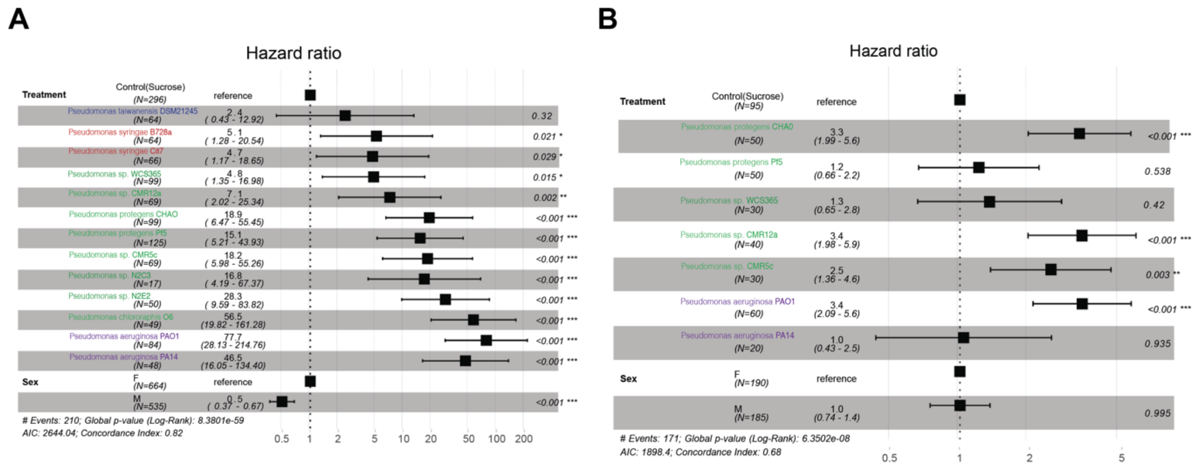


**Figure S3. Hazard ratios from Cox proportional hazards model.** Forest plots displaying the hazard ratio (HR) and 95% confidence intervals for each bacterial treatment. The significance level is indicated by asterisks. A) *Drosophila melanogaster* survival experiments. B) *Drosophila suzukii* survival experiments.

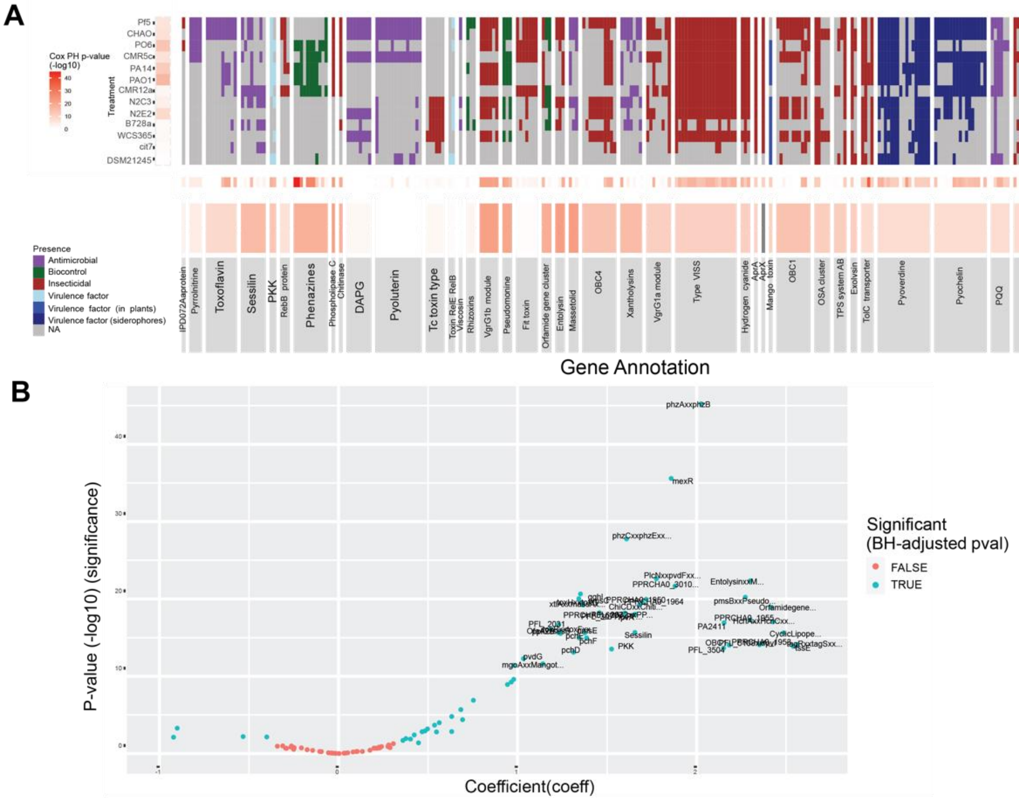


**Figure S4. Analysis of genes associated with *D. melanogaster* survival**

A) Cox proportional hazard and random forest models were used to predict the survival of *D. melanogaster* based on the presence or absence of specific genes. The p-values from the Cox proportional hazards model are represented by the red boxes, indicating the potential association between the presence of a particular gene and the risk of mortality in flies. The red boxes on the y-axis explain the p-value of survival probability after inoculation with each strain. The red boxes on the top horizontal row show the individual gene identity risk p-value, while the bottom horizontal row shows the p-value for gene annotation. The presence of genes is colour-coded based on their specific functions from the literature. B) Volcano plot of the genes based on p-value and coefficient. Significant genes are labeled in blue.

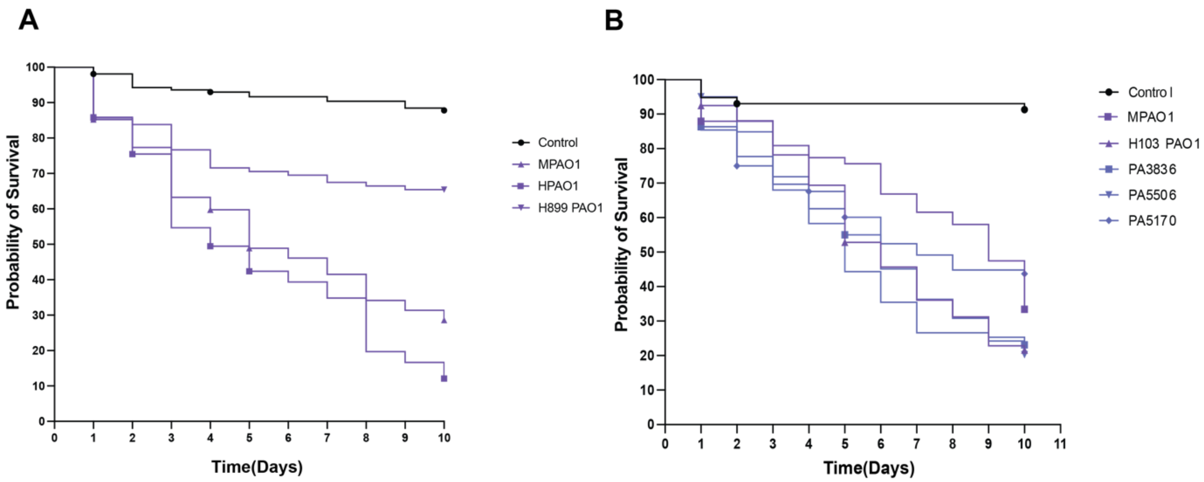


**Figure S5. Testing the virulence of *P. aeruginosa* parental strains.**
A) Kaplan-Meier (KM) survival curves of *D. melanogaster* Oregon-R flies following oral infection with *Pseudomonas* aeruginosa PA01 (MPAO1, H103PAO1 and H899PAO1) (OD600 = 100) or control 5% sucrose solution. B) Kaplan-Meier (KM) survival curves of *D. melanogaster* Oregon-R flies following oral infection with *Pseudomonas* aeruginosa PA01 (MPAO1 and H103PAO1), as well as the non-coding mutations PA3836, PA5506, PA5170, (OD600 = 100) or control 5% sucrose solution.

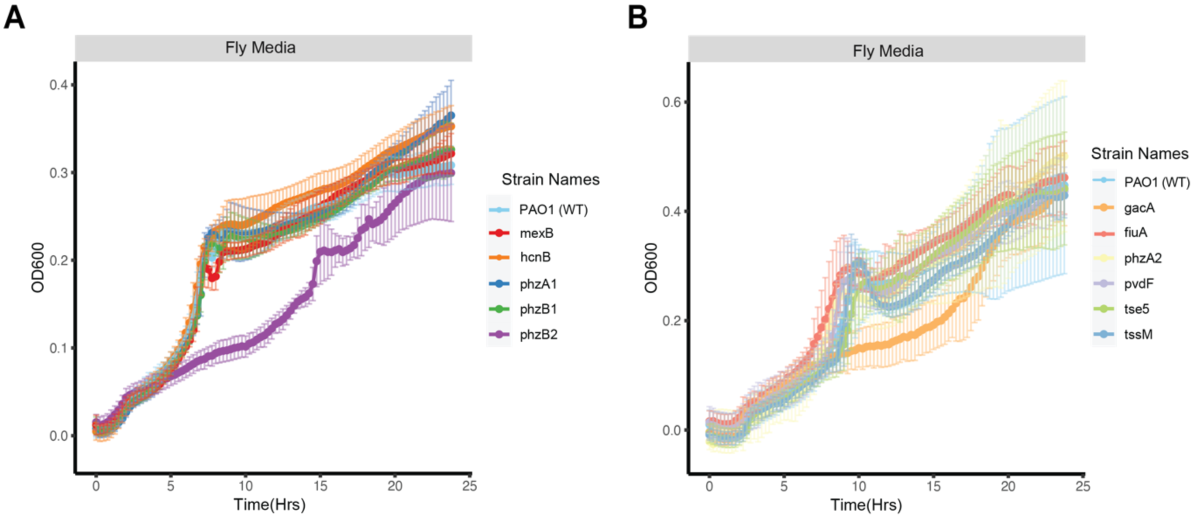


**Figure S6. Growth curves in M9 medium supplemented with fly extract.** Bacterial strains were grown in M9 medium supplemented with fly extract. A) Growth comparison using batch 1 fly extract, assessing PAO1 wild type against transposon mutants *mexB*, *hcnB*, *phzA1*, *phzB1*, and *phzB2*. B) Growth comparison using batch 2 fly extract, assessing PAO1 wild type against transposon mutants *gacA*, *fiuA*, *phzA2*, *pvdF*, *tse5*, and *tssM*.
